# Mapping metabolic oscillations during cell cycle progression

**DOI:** 10.1101/2020.01.31.928267

**Authors:** Irena Roci, Jeramie D. Watrous, Kim A. Lagerborg, Mohit Jain, Roland Nilsson

## Abstract

Proliferating cells must synthesize a wide variety of macromolecules while progressing through the cell cycle, but the coordination between cell cycle progression and cellular metabolism is still poorly understood. To identify metabolic processes that oscillate over the cell cycle, we performed comprehensive, non-targeted liquid chromatography-high resolution mass spectrometry (LC-HRMS) based metabolomics of HeLa cells isolated in the G_1_ and SG_2_M cell cycle phases, capturing thousands of diverse metabolite ions. When accounting for increased total metabolite abundance due to cell growth throughout the cell cycle, 18% of the observed LC-HRMS peaks were at least 2-fold different between the stages, consistent with broad metabolic remodeling throughout the cell cycle. While most amino acids, phospholipids, and total ribonucleotides were constant across cell cycle phases, consistent with the view that total macromolecule synthesis does not vary across the cell cycle, certain metabolites were oscillating. For example, ribonucleotides were highly phosphorylated in SG_2_M, indicating an increase in energy charge, and several phosphatidylinositols were more abundant in G_1_, possibly indicating altered membrane lipid signaling. Within carbohydrate metabolism, pentose phosphates and methylglyoxal metabolites were associated with the cycle. Interestingly, hundreds of yet uncharacterized metabolites similarly oscillated between cell cycle phases, suggesting previously unknown metabolic activities that may be synchronized with cell cycle progression, providing an important resource for future studies.

## Introduction

The cell cycle is a fundamental biological process that requires close coordination of complex cellular activities to enable copying of internal structures and division into two daughter cells. While much research has been devoted to the signaling events that regulate progression through the cell cycle, the interplay between the cell cycle and the underlying metabolic machinery remains poorly understood. It is clear that cells must activate a multitude of anabolic processes to support biosynthesis of all their components, while catabolism of certain nutrients must also increase, since biosynthesis requires large amounts of energy. Cells must coordinate all these processes while ensuring timely synthesis of various cellular structures, suggesting that certain aspects of metabolism should be coordinated with cell cycle progression. Besides its fundamental importance in cell biology, the fact that many human disorders involve cell proliferation also renders the cell cycle important for biomedicine. Metabolic enzymes are often good drug targets (1), and several commonly used anti-proliferative drugs target the few well-known cell cycle-associated metabolic pathways, notably synthesis of deoxynucleotides (antifolates, thymidylate synthesis inhibitors) (2) and polyamines (3). Uncovering cell cycle-associated metabolic processes is therefore of interest to find new avenues towards controlling cell proliferation.

A number of observations suggest that some aspects of cellular metabolism are indeed coordinated with cell cycle progression. The most obvious is perhaps the synthesis of nuclear DNA in S phase, which coincides with a burst of deoxyribonucleotide triphosphate (dNTP) synthesis (4). Enzymes in polyamine metabolism have also been found to increase during the S-G_2_ phase (5), and may be related to chromatin duplication and rearrangements. RNA synthesis on the other hand appears to be continuous during the G_1_-S-G_2_ phases of the cycle, but ceases during M phase when chromatin is condensed (6), indicating that *de novo* ribonucleotide (NTP) synthesis (which constitutes the majority of cell nucleotides) is regulated differently than dNTP synthesis. Within energy metabolism, cell cycle-associated utilization of glycogen (7), pentose phosphate pathway activity (8), and fluxes in the TCA cycle (9) have been observed to be associated with cell cycle phase in various model systems. On the other hand, cell biomass appears to grow continuously throughout the cell cycle (10), suggesting that protein and membrane synthesis, and hence amino acid and lipid demand, should not vary much over the cell cycle, although the precise growth dynamics are still a matter of debate (11).

Beyond these particular findings, there are few systematic studies of metabolites associated with the cell cycle. This is partly because it is difficult to obtain pure cell populations at a given cell cycle phase without disturbing metabolism with synchronization reagents. For some cell types, cell cycle progression can be synchronized by blocking key steps such as nucleotide synthesis or nuclear spindle formation, but such synchronization methods can themselves alter metabolic processes, and many human cell types are not amenable to these techniques. To address this problem, we have developed a methodology for metabolomics analysis of cells separated by cell sorting (12,13). This approach avoids artefacts introduced by synchronization agents, such as transient responses that are unrelated to true cycling behavior (14), and allows directly observing the metabolic state at particular stages of the cell cycle, as they naturally occur in undisturbed cell cultures. Here, we used this technique to chart the abundance of metabolites in the G_1_ or SG_2_M cell cycle phases using untargeted high-resolution LC-HRMS, thus providing a resource for future studies of cell-cycle associated metabolic pathways.

## Results and Discussion

### A resource of cell cycle-associated metabolites

To identify metabolites associated with a particular cell cycle phase, we performed sorting of cultured HeLa cells based on DNA content as quantified by Hoechst-34580, resulting in 2n and 4n fractions, as previously described (13). We have previously demonstrated that these fractions contain highly pure populations of G_1_ and SG_2_M cells (13), and consequently we refer to these sorted populations as G_1_ and SG_2_M in what follows. We then analyzed metabolite abundances in 80,000 HeLa cells from G_1_ and SG_2_M using liquid chromatography-high resolution mass spectrometry (LC-HRMS). In sorted HeLa cells, we detected 3,426 LCMS peaks representing potential metabolites, of which 1,549 could be annotated with at least one putative metabolite identity based on exact mass, matched against the Human Metabolite Database (15). Of these, 921 peaks passed manual quality checks (Figure 1a), and were considered for further analysis. To assess the quality of these annotations, we selected 268 peaks whose identity was known from pure standards, and found that all these identities were also found in HMDB, suggesting that our putative metabolite annotations are of good quality. In addition, by collecting MS/MS spectra from the observed LCMS peaks, we were able to annotate another 163 peaks with metabolite identities, and these MS/MS annotations were consistent with the annotations from pure standards. The full data set is available online (**Supplementary Table 1**) as a resource for the community.

**Figure 1.**
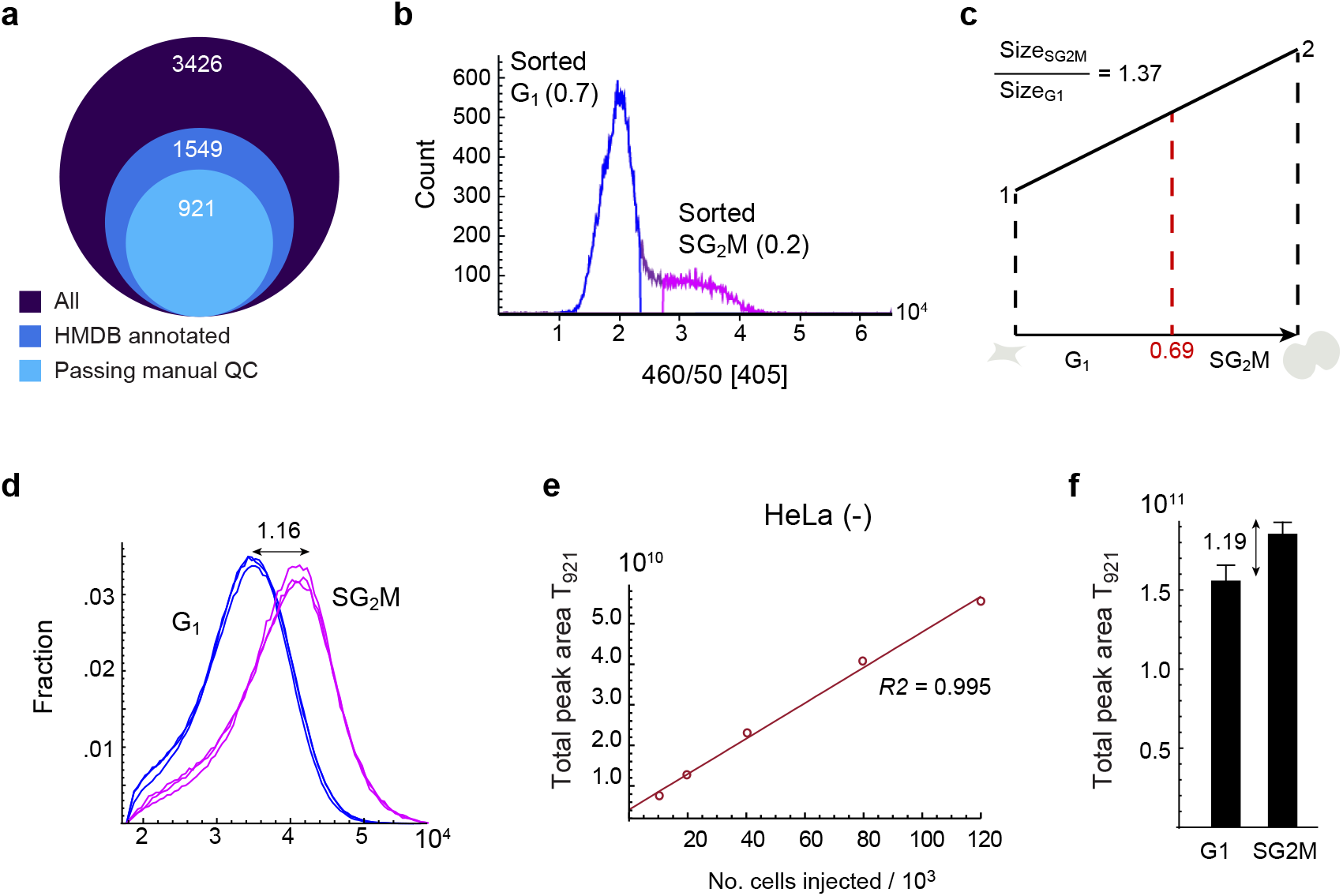
Metabolite content as a normalization factor for cell cycle progression. a) Diagram of metabolite peak numbers. 3426 peaks were identified from untargeted analysis, 1549 peaks were annotated from HMDB, and 921 peaks were manually curated. b) DNA histogram of sorted cells. Fraction of sorted G_1_ and SG_2_M phase cells is indicated in the plot. c) Illustration of theoretical cell growth during the cell cycle. When the fraction of G_1_ cells is 0.69, the calculated theoretical size ratio between SG_2_M vs G_1_ is 1.37. d) Forward scatter of cells in G_1_ and SG_2_M phases as a measure of cell size. The ratio of the median for SG_2_M vs G_1_ is 1.16. e) The correlation of total peak area of 921 (T_921_) metabolites with the number of injected cells. f) T_921_ of cells in G_1_ and SG_2_M phases, respectively. T_921_ ratio of SG_2_M vs G_1_ is 1.19, and is used to normalize metabolite data.

### Metabolite content as a normalization factor for cell growth

An important initial question was how metabolite abundance data should be normalized, given that cell biomass increases over the cell cycle, and metabolite abundance is probably best considered relative to cell size. We therefore set out to estimate the relative cell size of an “average” cell in the G_1_ and SG_2_M phases, respectively. From the DNA histograms obtained during the cell sorting, we estimated the fraction of G_1_ and SG_2_M cells to be 0.7 and 0.2, respectively (Figure 1b). Based on the steady-state distribution of cycling cells (16), we then estimated the fractional length of the G_1_ and SG_2_M phases to be 0.69 and 0.31, respectively, which gives a theoretical cell size ratio of 1.37 between the G_1_ and SG_2_M fractions, assuming linear growth (Figure 1c). On the other hand, using cytometry forward scatter as a proxy for cell size (17), we estimated a SG_2_M/G_1_ ratio of 1.16 (Figure 1d). To more directly estimate the difference in metabolite content between sorted fractions, we first established that the total peak area of the curated 921 (*T*_921_) peaks closely corresponds to cell number (Figure 1e), and hence should reflect biomass. In sorted cells, the SG_2_M/G_1_ ratio of *T*_921_ was about 1.19 (Figure 1f), which agrees well with the forward scatter estimates, indicating that total metabolite content correlates with cell size. To compensate for this factor, we express all metabolite peak abundance data as SG_2_M/G_1_ ratios normalized to *T*_921_, to reflect relative intracellular concentrations as the cell increases in volume during cell cycle, rather than amounts per cell.

### Oscillating and housekeeping central metabolism in cycling cells

The majority of LCMS peaks (82%) did not differ in relative abundance between phases, while 74 (7%) were >2-fold higher in G_1_, and 107 (11%) were >2-fold higher in SG_2_M (Figure 2a). Interestingly, this is similar to the 15–18% of mRNA transcripts and proteins that are estimated to be cyclic in HeLa cells (18). Although our data only captures a subset of polar metabolites, this data suggests that most metabolic processes are independent of cell cycle phase, while a smaller set of pathways or enzymes are oscillating. This seems reasonable, given that cell biomass grows continuously over the cell cycle (19), so that most metabolic products should be in constant demand, and the corresponding metabolites should be roughly constant when normalized to cell size. Reassuringly, we observed that dNTPs, which are known to be synthesized only during S phase (20,21), were exclusively found in the SG_2_M fraction (Figure 2b), as previously reported (13). We also observed that deoxynucleotide monophosphates (dNMPs), which are intermediates in the dNTP salvage pathway, were several-fold more abundant in the SG_2_M phases (Figure 2c). This may reflect ongoing salvage of dNTPs derived from degraded DNA (22), or from DNA repair mechanisms (23), which are highly active during S phase in transformed cells.

**Figure 2.**
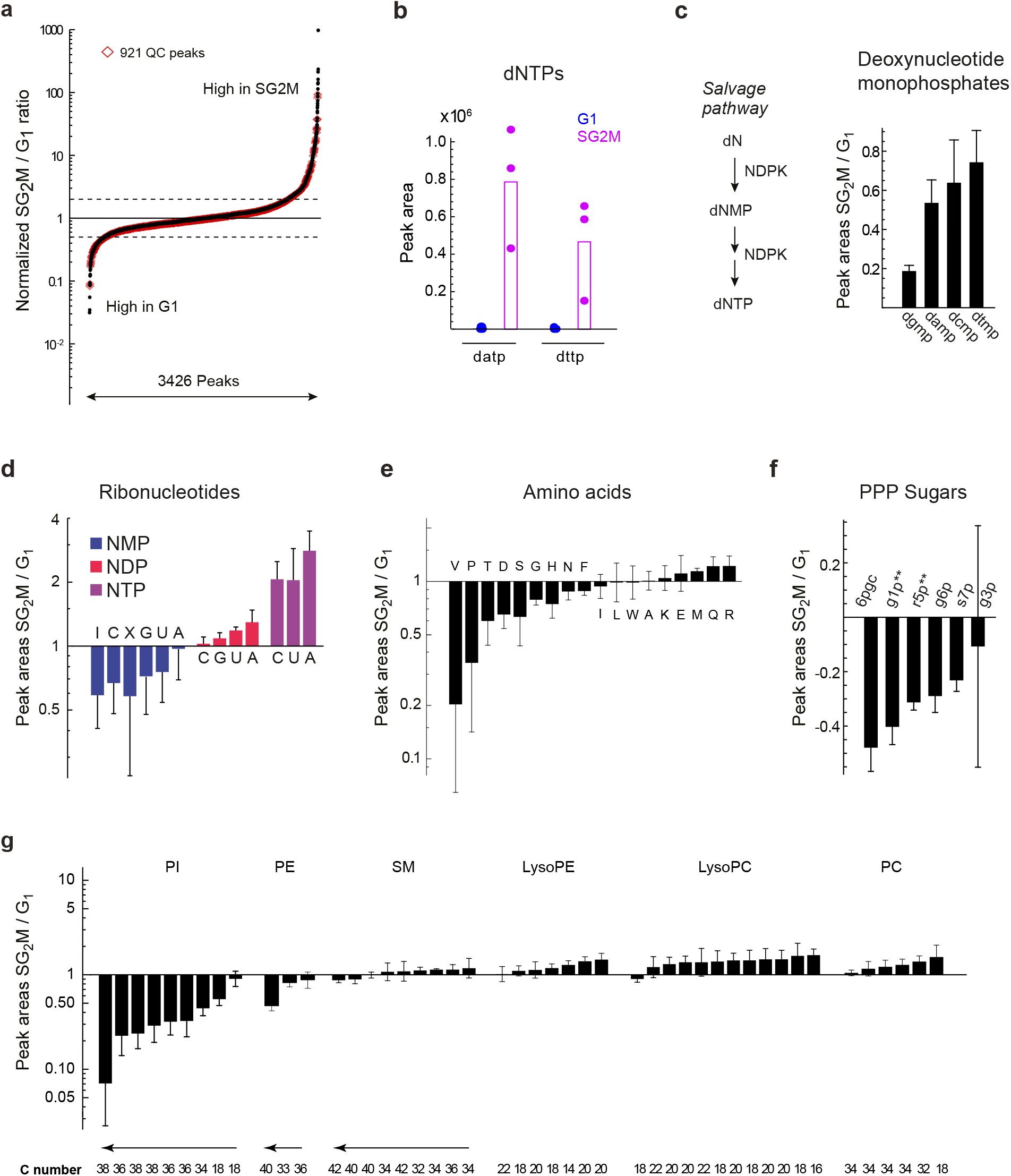
Oscillating and housekeeping metabolites in cycling cells. a) S-plot of metabolite relative abundance ratios in SG_2_M vs G_1_ phase cells. b) Peak areas of (deoxyadenosine triphosphate) datp and (deoxythymine triphosphate) dttp in G_1_ (blue) and SG_2_M (red) cells. (c-g) Relative abundances ratios (SG_2_M vs G_1_) for: c) Deoxyri-bonucleotide monophosphates, d) Ribonucleotides, e) Amino acids, f) PPP sugars, g) phospholipid subtypes. All ratios of peak areas presented in this figure are calculated as Log10 of peak areas in SG_2_M vs G_1_, and plotted as sorted based on mean of ratios SG_2_M/G_1_.

While total ribonucleotides (NTPs) did not vary between the cell cycle phases, the ribonucleotide triphosphates ATP, CTP and UTP (GTP was not measurable) were all between 3.5 to 5-fold higher in the SG_2_M phases, while the corresponding diphosphates were comparable between phases, and the monophosphates were lower (Figure 2d). These differences indicate a higher energy charge (24) in SG_2_M phase cells, suggesting that energy metabolism oscillates over the cell cycle. Oscillation in ATP level was previously reported in synchronized mouse 3T3 fibroblasts (25), and less pronounced oscillation in GTP level in HL-60 cells (26) and in yeast (27), suggesting that the phenomenon is deeply conserved. The nucleoside diphosphokinase enzyme (NDP kinase, encoded by *NME1–9* in humans), which synthesizes NTPs from NDPs, was also found to continuously increase in activity during the cell cycle of fission yeast, with highest peak in S phase (28), possibly explaining the simultaneous increase in phosphorylation state of all nucleotides. Oscillating energy charge might be related to increased demand for NTPs in biosynthetic processes (29) in the SG_2_M phases, for example during DNA replication, or reflect changes in ATP synthesis. In this regard, glycolysis and TCA cycle metabolites did not differ markedly, but acetyl-CoA was almost 10-fold higher in SG_2_M (Supplementary Table S2). On the other hand, some phosphate sugars in the pentose phosphate pathway (PPP), which provides ribose sugars and NADPH, were more abundant in G_1_ phase (Figure 2e), consistent with previous reports using ^13^C labeling (13).

As protein synthesis is constantly ongoing during G_1_–S–G_2_ phases, we expected abundances of the proteinogenic amino acids to be mostly independent of cell cycle. Indeed, most amino acids did not oscillate; however, valine, but not the other branched-chain amino acids, stood out as 10-fold higher in G_1_ phase (Figure 2f). Interestingly, some carcinoma cells have been found to arrest in G_1_ phase when valine is removed from the medium (30), suggesting a specialized role for the amino acid in this cell cycle phase. Arginine was somewhat higher in SG_2_M, in line with our previous report (13). The phosphatidylcholines, which are major structural lipids in cell membranes, were generally constant across the cell cycle. However, phosphatidylinositol peaks (PIs) were in higher abundance in G_1_ (Figure 2g). PI signaling events at the plasma membrane are known to be important for transition through G_1_ phase (31), and certain PIs have also been found in the nucleus, where they are more abundant during G_1_ (32,33). An interesting hypothesis is that PI content of membranes might be dynamically regulated to facilitate PI signaling in specific cell cycle phases. Overall, these observations support the notion that major biosynthetic processes are constant across the cell cycle, while specific metabolites or pathways are synchronized with it.

### Identification of lactoylglutathione, a novel cell-cycle associated metabolite

The majority of the cell cycle-associated LCMS peaks did not have a clear annotation. We expect that a large fraction of these peaks are either in-source fragments or natural isotopomers of already annotated metabolites, as such features are common in LCMS (34). However, we also found a number of putative metabolites that have not previously been associated with the cell cycle. For example, a peak at m/z = 380.112 was putatively annotated as S-Lactoylglutathione, and was about 3-fold more abundant in G_1_ than in SG_2_M (Figure 3a). S-Lactoylglutathione is an intermediate in the conversion of methylglyoxal to pyruvate (Figure 3b), and increased levels of this metabolite could reflect altered levels of methylglyoxal between G_1_ and SG_2_M. To ensure that this peak did not result from in-source fragmentation, we computed all possible parent-fragment pairs among the 921 peaks, and verified that no other peak was a possible parent (Supplementary Table S3). To support the identity of this peak (RT 10.6-11.2 min), we obtained the MS/MS spectrum of S-Lactoylglutathione from sorted cells, and confirmed that it is almost identical with MS/MS data of S-Lactoylglutathione in a mix of pure standards, supporting the identity of this peak (Supplementary Fig. S1). In addition, based on matching MS/MS fragments with Metlin database (35), some of the fragments (m/z 76.0223, 84.0451, 233.059) matched glutathione fragments. To further investigate methylglyoxal metabolism in cycling cells, we performed follow-up isotope tracing experiments using ^13^C_5_-glutamine (13), which should yield a ^13^C_5_ glutathione moiety, and with ^13^C_1_-glucose (36), which is expected to label the lactoyl moiety as 50% ^13^C_5_. Indeed, we observed these mass isotopomers in the S-Lactoylglutathione peak (Figure 3c), demonstrating that the compound was endogenously synthesized. The S-Lactoylglutathione peak was not found in fresh medium (data not shown), indicating that it is an intracellular metabolite. A glutathione peak and a putative methylglyoxal peak were also detected in these samples (Figure 3d), and were also somewhat more abundant in G_1_. These results suggest that methylglyoxal metabolism may be cell-cycle associated, consistent with reports that methylglyoxal treatment of cells can cause cell cycle arrest (37–39).

**Figure 3.**
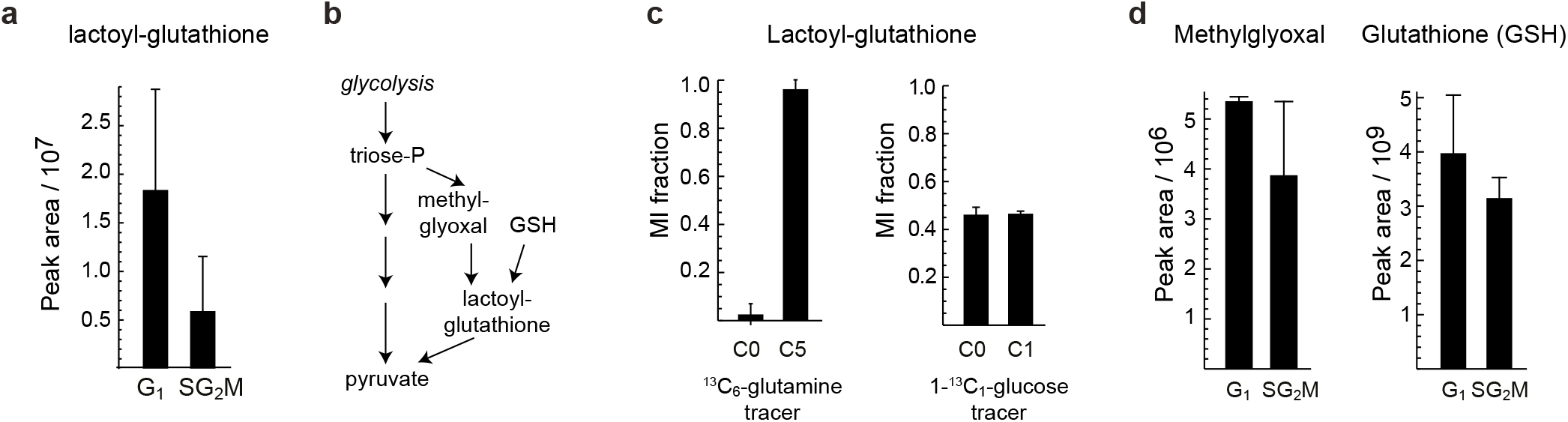
Identification of unknown cycling metabolites. a) Relative abundance of peak 223, annotated as S-Lactoylglutathione. b) Methylglyoxal detoxification reactions where S-Lactoylglutathione is involved. c) Labeling of S-Lactoylglutathione in ^13^C_6_-glutamine (HeLA) and 1-^13^C_1_-glucose (HCT116) tracing experiments. Metabolite abundance ratios are calculated as Log10 of peak areas in SG_2_M vs G_1_. d) Relative abundance of methylglyoxal and reduced glutathione in sorted samples.

## Conclusions

This study provides an overview of polar metabolites associated with cell cycle phases in HeLa cells, which we hope will serve as a valuable resource for investigating the coordination between metabolic processes and cell cycle progression. The data presented here is orthogonal to existing studies based on synchronization methods (9), providing direct observations of metabolite abundances in cell cycle phases of undisturbed cycling cells in culture. We have shown in a series of examples that multiple metabolic pathways exhibit such coordination, of which many are yet to be studied in depth. One potential application is the identification of cell-cycle associated enzymes, which might be exploited as targets for anti-proliferative drugs. Also, our data spotlight several putative metabolites that appear closely associated with the G_1_ phase, for which there are currently no good markers.

There are several limitations that must be acknowledged. First, the cell sorting procedure clearly affects measured peak areas for a variety of metabolites (12). It is still unclear to what extent this reflects actual distortion of metabolism during cell sorting, and to what extent the effect is merely due to a difference in the chemical matrix of the sample (which affects LC-HRMS peak intensities). To mitigate bias due to cell sorting, all comparisons were made in pairwise fashion between G_1_ and SG_2_M populations obtained within the same cell sorting experiment, which presumably were affected in the same way. Nevertheless, it is important to validate results of interest in follow-up experiments with orthogonal techniques, such as chemical synchronization or elutriation methods. In addition, it should be emphasized that the metabolite identities reported in this study are tentative, and in most cases require validation experiments for confirmation, prior to other follow-up experiments to further investigate the role of the associated metabolic pathways in cell cycle progression. Yet, as our data shows, the putative identities provided by matching by exact mass to HMDB often contain the correct identity, although there may also be additional false annotations due to isobaric species. In the S-lactoylglutathione case that we chose for further validation, our experimental data clearly supports the annotation. We hope that these results and the data sets provided by this study will be a valuable resource and inspire further research in this field.

## Methods

### Cell culture and sorting

HeLa (human cervix adenocarcinoma) cells were cultured for 48 hours in RPMI-1640 (Gibco, 61870-010) + 5% FBS (Gibco, 16140-071) + 1% Pen-Strep (Gibco, 15140-122). Prior to sorting, cells were incubated for 15 minutes in Hoechst staining solution in dark at 37°C. The Hoechst staining solution was prepared by adding 11μL Hoechst-34580 (Sigma Aldrich, 63493) stock (1mg/mL in dH_2_O) and 10μL Verapamil (Sigma Aldrich, PHR1131) stock (1mg/mL in DMSO) to 10mL warm culture medium.

At the end of the incubation time, culture medium was aspirated, cells were rinsed with PBS, treated with trypsin for 5 minutes at 37°C, and transferred to a new tube. Culture plates were rinsed with RPMI-1640 + 5% FBS to block trypsin, and solution was added to a respective tube. Then, cell solutions were centrifuged at 1500rpm for 3 minutes, and supernatant was aspirated. The cell pellet was resuspended in PBS + 5% dFBS at a concentration of 1×10^6^ cells per mL, filtered through Flowmi cell strainers (Bel-Art Products, 734-5950), and transferred to a 5mL polyprolylene tube (BD Falcon, 352063).

Cells in suspension were sorted into 5mL Eppendorf tubes (Thermo Fischer, 0030119380) based on their DNA staining into Hoechst negative (2n, G_1_) and Hoechst positive (4n, SG_2_M), at an event rate of 1000 events/second using 100μm nozzle at 4°C.

### Metabolite extraction and LC-HRMS analysis

After the sorting procedure, sorted fractions were centrifuged at 1500rpm for 3 minutes at 4°C. The supernatant was aspirated, and pellet was quickly resuspended with 5μL dH2O. 500μL dry ice cold methanol was added to the cell suspension to quench and extract metabolites. The cell extracts were transferred to a new tube and kept in −80°C until mass spectrometry analysis. Sample analysis using Liquid Chromatography - High Resolution Mass Spectrometry (LC-HRMS) was performed as previously described (13).

### LC-HRMS Data processing

#### Targeted analysis

LC-HRMS chromatograms were retrieved from raw instrument data files using the data access layer mzAccess (40), and processed using Wolfram Mathematica v.10 (Wolfram Research). Metabolites from main classes of amino acids, ribonucleotides, deoxyribonucleotides and pentose phosphate pathway (PPP) sugars were identified based on m/z and retention time (RT) data obtained from pure standards on the same LCMS method.

#### Untargeted analysis

Untargeted analysis was implemented using an in-house developed peak detection software, with the prerequisite that detected peaks have a highest point intensity of at least 500,000 and be present in at least 3 out of 6 samples (3 x G_1_ and 3 x SG_2_M). After excluding peaks that were present in blanks, a list of 3426 peaks was obtained. Based on the estimated m/z within 10 ppm accuracy and accounting for +H and −H adducts the list was matched with peak entries from Human Metabolome Database (HMDB) (15), and 1549 peaks were annotated. Other annotations were retrieved from matching collected MS/MS spectra against the public library GNPS online tool (41). The putative metabolite annotations, m/z, RT, HMDB annotations and GNPS annotations are available in Supplementary Table 1.

#### Normalization factor

To compensate for growth and biomass increase during cell cycle progression we calculated total metabolite peak area of 921 metabolites, and used the ratio of SG_2_M to G_1_ (1.19) as a normalization factor for peak area calculations.

## Acknowledgements

This work was supported by grants from the Strategic Research Programme in Cancer at Karolinska Institutet (I.R., R.N.); and the Foundation for Strategic Research (FFL12-0220: I.R., R.N.); Robert Lundberg Memorial Foundation (2017-00516: IR); the UC San Diego Frontiers of Innovation Scholars Program (K.A.L.), as well as grants from the National Institutes of Health (S10OD020025, R03H133720 and R01ES027595 to M.J.,and K01DK116917 to J.D.W.).

## Author contributions

R.N., I.R., M.J. designed research; I.R. performed sorting, flow cytometry, microscopy and knockdown experiments; J.D.W. and K.A.L. processed and analyzed samples in LC-HRMS; IR and R.N. analyzed data; R.N. and I.R. wrote the manuscript.

## SUPPLEMENTARY FIGURES

**Figure S1.**
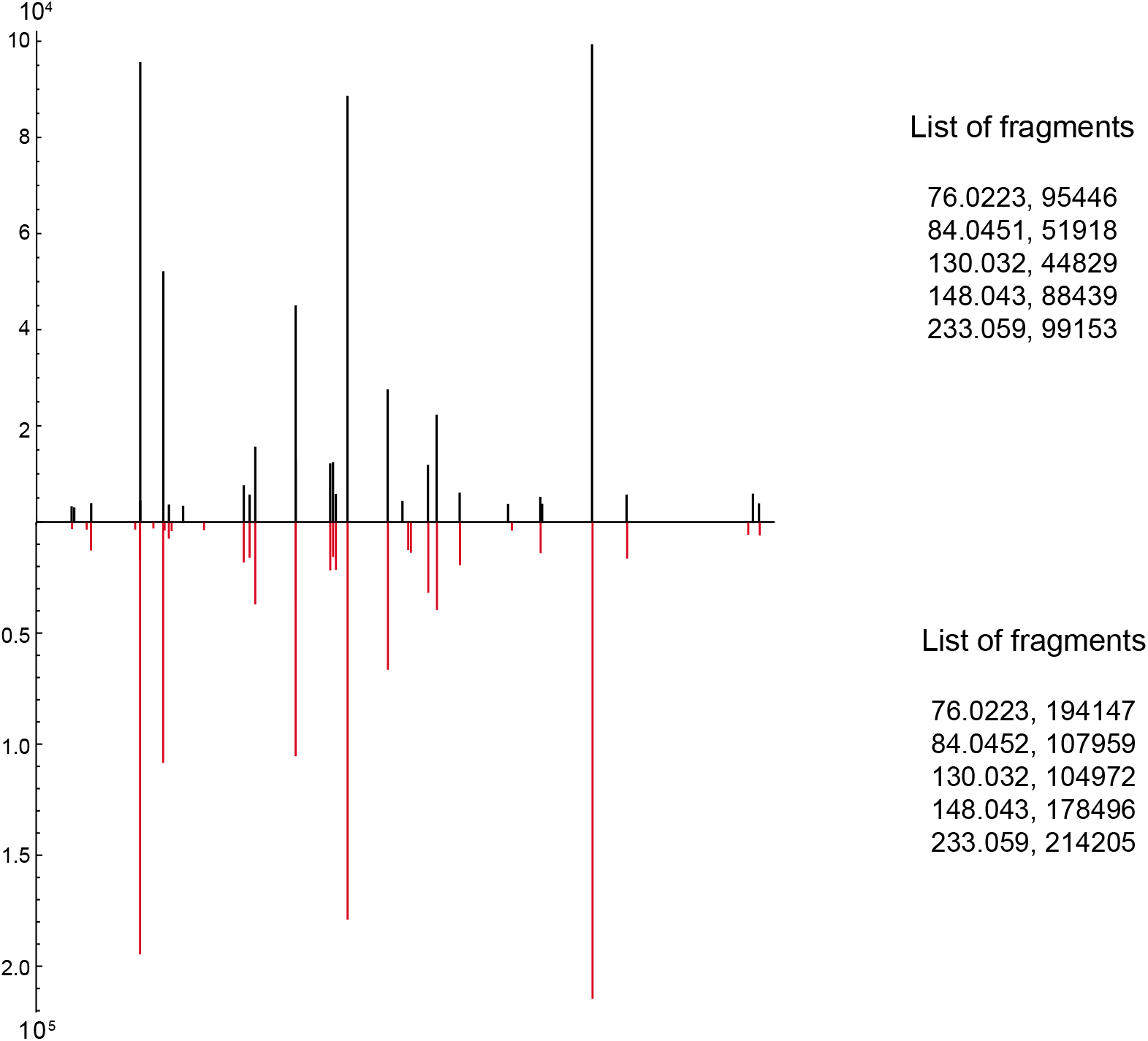
MS/MS data for peak 223 (S-Lactoylglutathione) in sorted samples (black) and standard mix (red).

## References

1. Hopkins AL, Groom CR 2002 The druggable genome. Nat Rev Drug Discov. 1:727–30.

2. Wilson PM, Danenberg P V., Johnston PG, Lenz H-J, Ladner RD 2014 Standing the test of time: targeting thymidylate biosynthesis in cancer therapy. Nat Rev Clin Oncol. Nature Publishing Group; 11:282–98.

3. Kim DJ, Roh E, Lee M-H, Oi N, Lim DY, Kim MO, et al. 2016 Herbacetin is a novel allosteric inhibitor of ornithine decarboxylase with antitumor activity. Cancer Res. NIH Public Access; 76:1146.

4. Reichard P 1988 Interactions between deoxyribonucleotide and DNA synthesis. Ann Rev Biochem. 57:349–74.

5. Fredlund JO, Johansson MC, Dahlberg E, Oredsson SM 1995 Ornithine decarboxylase and S-adenosylmethionine decarboxylase expression during the cell cycle of Chinese hamster ovary cells. Exp Cell Res. 216:86–92.

6. Taylor JH 1960 Nucleic acid synthesis in relation to the cell division cycle. Ann N Y Acad Sci. 90:409–21.

7. Rousset M, Chevalier G, Rousset J-P, Dussaulx E, Zweibaum A 1979 Presence and cell growth-related variations of glycogen in human colorectal adenocarcinoma cell lines in culture. Cancer Res. 39:531–4.

8. Vizán P, Alcarraz-Vizán G, Díaz-Moralli S, Solovjeva ON, Frederiks WM, Cascante M 2009 Modulation of pentose phosphate pathway during cell cycle progression in human colon adenocarcinoma cell line HT29. Int J Cancer. Wiley Subscription Services, Inc., A Wiley Company; 124:2789–96.

9. Ahn E, Kumar P, Mukha D, Tzur A, Shlomi T 2017 Temporal fluxomics reveals oscillations in TCA cycle flux throughout the mammalian cell cycle. Mol Syst Biol. 13.

10. Mitchison JM 2003 Growth During the Cell Cycle. Int Rev Cytol. 226:165–258.

11. Ginzberg MB, Kafri R, Kirschner M 2015 On being the right (cell) size. Science (80-). 348:1245075-1–7.

12. Roci I, Gallart-Ayala H, Schmidt A, Watrous J, Jain M, Wheelock CE, et al. 2016 Metabolite Profiling and Stable Isotope Tracing in Sorted Subpopulations of Mammalian Cells. Anal Chem. American Chemical Society; 88:2707–13.

13. Roci I, Watrous JD, Lagerborg KA, Lafranchi L, Lindqvist A, Jain M, et al. 2019 Mapping Metabolic Events in the Cancer Cell Cycle Reveals Arginine Catabolism in the Committed SG2M Phase. Cell Rep. 26:1691–1700.e5.

14. Whitfield ML, Sherlock G, Saldanha AJ, Murray JI, Ball CA, Alexander KE, et al. 2002 Identification of genes periodically expressed in the human cell cycle and their expression in tumors. Mol Biol Cell. 13:1977–2000.

15. Wishart DS, Jewison T, Guo AC, Wilson M, Knox C, Liu Y, et al. 2013 HMDB 3.0--The Human Metabolome Database in 2013. Nucleic Acids Res. 41:D801–7.

16. Toettcher JE, Loewer A, Ostheimer GJ, Yaffe MB, Tidor B, Lahav G 2009 Distinct mechanisms act in concert to mediate cell cycle arrest. Proc Natl Acad Sci. 106:785–90.

17. Tzur A, Moore JK, Jorgensen P, Shapiro HM, Kirschner MW 2011 Optimizing optical flow cytometry for cell volume-based sorting and analysis. PLoS One. Public Library of Science; 6:e16053.

18. Grant GD, Brooks L, Zhang X, Mahoney JM, Martyanov V, Wood TA, et al. 2013 Identification of cell cycle-regulated genes periodically expressed in U2OS cells and their regulation by FOXM1 and E2F transcription factors. Mol Biol Cell. 24:3634–50.

19. Son S, Tzur A, Weng Y, Jorgensen P, Kim J, Kirschner MW, et al. 2012 Direct observation of mammalian cell growth and size regulation. Nat Methods. Nature Research; 9:910–2.

20. Bray G, Brent TP 1972 Deoxyribonucleoside 5’-triphosphate pool fluctuations during the mammalian cell cycle. Biochim Biophys Acta. 269:184–91.

21. Skoog KL, Nordenskjold BA, Bjursell KG 1973 Deoxyribonucleoside-Triphosphate Pools and DNA Synthesis in Synchronized Hamster Cells. Eur J Biochem. Blackwell Publishing Ltd; 33:428–32.

22. Fasullo M, Endres L 2015 Nucleotide salvage deficiencies, DNA damage and neurodegeneration. Int J Mol Sci. Multidisciplinary Digital Publishing Institute (MDPI); 16:9431–49.

23. Berdis AJ 2009 Mechanisms of DNA Polymerases. Chem Rev. 109:2862–79.

24. Atkinson DE 1968 Energy charge of the adenylate pool as a regulatory parameter. Interaction with feedback modifiers. Biochemistry. American Chemical Society; 7:4030–4.

25. Marcussen M, Larsen PJ 1996 Cell cycle-dependent regulation of cellular ATP concentration, and depolymerization of the interphase microtubular network induced by elevated cellular ATP concentration in whole fibroblasts. Cell Motil Cytoskeleton. 35:94–9.

26. Szekeres T, Fritzer M, Pillwein K, Felzmann T, Chiba P 1992 Cell cycle dependent regulation of IMP dehydrogenase activity and effect of tiazofurin. Life Sci. 51:1309–15.

27. Thomas KC, Dawson PS 1977 Variations in the adenylate energy charge during phased growth (cell cycle) of Candida utilis under energy excess and energy-limiting growth conditions. J Bacteriol. 132:36–43.

28. Dickinson JR 1983 Nucleoside diphosphokinase and cell cycle control in the fission yeast Schizosaccharomyces pombe. J Cell Sci. 60:355–65.

29. Atkinson DE (1977.) Interactions between Regulatory Parameters. Cell Energy Metab its Regul. Elsevier; p. 175–200.

30. Sun X, Zhang N, Li K, Liu M, Zhi X, Jiang X, et al. 2003 Branched chain amino acid imbalance selectively inhibits the growth of gastric carcinoma cells in vitro. Nutr Res. Elsevier; 23:1279–90.

31. Jones SM, Kazlauskas A 2001 Growth-factor-dependent mitogenesis requires two distinct phases of signalling. Nat Cell Biol. Nature Publishing Group; 3:165–72.

32. Clarke JH, Letcher AJ, D’santos CS, Halstead JR, Irvine RF, Divecha N 2001 Inositol lipids are regulated during cell cycle progression in the nuclei of murine erythroleukaemia cells. Biochem J. 357:905–10.

33. York JD, Majerus PW 1994 Nuclear Phosphatidylinositols Decrease during S-phase of the Cell Cycle in HeLa Cells*. J BIOWCICAL Chem. 269:7847–50.

34. Mahieu NG, Patti GJ 2017 Systems-Level Annotation of a Metabolomics Data Set Reduces 25 000 Features to Fewer than 1000 Unique Metabolites. Anal Chem. American Chemical Society; 89:10397–406.

35. Guijas C, Montenegro-Burke JR, Domingo-Almenara X, Palermo A, Warth B, Hermann G, et al. 2018 METLIN: A Technology Platform for Identifying Knowns and Unknowns. Anal Chem. 90:3156–64.

36. Grankvist N, Watrous JD, Lagerborg KA, Lyutvinskiy Y, Jain M, Nilsson R 2018 Profiling the Metabolism of Human Cells by Deep 13C Labeling. Cell Chem Biol. 25:1419–1427.e4.

37. Kang Y, Edwards LG, Thornalley PJ 1996 Effect of methylglyoxal on human leukaemia 60 cell growth: Modification of DNA, G1 growth arrest and induction of apoptosis. Leuk Res. 20:397–405.

38. Kani S, Nakayama E, Yoda A, Onishi N, Sougawa N, Hazaka Y, et al. 2007 Chk2 kinase is required for methylglyoxal-induced G 2 /M cell-cycle checkpoint arrest: implication of cell-cycle checkpoint regulation in diabetic oxidative stress signaling. Genes to Cells. 12:919–28.

39. Braun JD, Pastene DO, Breedijk A, Rodriguez A, Hofmann BB, Sticht C, et al. 2019 Methylglyoxal down-regulates the expression of cell cycle associated genes and activates the p53 pathway in human umbilical vein endothelial cells. Sci Rep. 9:1152.

40. Lyutvinskiy Y, Watrous JD, Jain M, Nilsson R 2017 A Web Service Framework for Interactive Analysis of Metabolomics Data. Anal Chem. 89:5713–8.

41. Wang M, Carver JJ, Phelan V V., Sanchez LM, Garg N, Peng Y, et al. 2016 Sharing and community curation of mass spectrometry data with Global Natural Products Social Molecular Networking. Nat Biotechnol. 34:828–37.

